# Dietary fiber-type-specific gut microbiome preconditioning drives differential susceptibility to sporadic colon cancer development

**DOI:** 10.64898/2026.09.21.753221

**Authors:** Sangshan Tian, Sumudu Rajakaruna, Gopi Yalavarthi, Umesh K Goand, Tinghua Chen, Won Gu, Justin Silverman, Fuhua Hao, Seth R. Bordenstein, Jordan E. Bisanz, Amit K. Tiwari, Andrew D. Patterson, Vishal Singh

## Abstract

Dietary fiber intake is broadly associated with reduced colorectal cancer (CRC) risk, yet processed fiber supplements exert inconsistent and occasionally opposing effects on colon tumorigenesis. How structurally distinct dietary fibers differentially shape the gut microbiome–metabolome axis to influence carcinogenic susceptibility remains poorly understood. We compared structurally distinct dietary fibers commonly added to processed foods in an experimental model of sporadic CRC. Four-week-old male mice were maintained on low-fiber control or fiber-supplemented diets (7.5%w/w: cellulose, agar, pectin, or inulin) for 28 weeks and received eight intraperitoneal azoxymethane injections (AOM; 7.5mg/kg). Temporal fecal microbiomes were profiled by 16S-rRNA sequencing, cecal metabolites quantified by ¹H-NMR spectroscopy, and microbiome–metabolome interactions assessed by co-occurrence networks. Refined inulin supplementation markedly elevated colon tumor incidence (∼70%) compared with all other groups (∼10–20%), and cecal extracts from inulin-fed mice promoted HT29 cancer-cell proliferation. Longitudinal microbiome profiling identified 36 inulin-specific microbial signatures converging on two axes: depletion of SCFA-producing and fermentation cross-feeding taxa and reprogramming of amino acid and nitrogen metabolism. Metabolomics confirmed succinate, fumarate, and lactate accumulation alongside propionate and amino acid depletion exclusively under AOM conditions, with no differences in carcinogen-naive animals. Two stable co-occurrence hubs were identified exclusively in AOM-treated animals, structurally integrating both functional axes. These findings demonstrate that inulin supplementation preconditions the gut microbiome toward a latent metabolic vulnerability, disrupted fermentation throughput and reprogrammed amino acid metabolism, that carcinogen exposure amplifies into a tumor-promoting ecosystem, establishing fiber structure as a critical and underappreciated determinant of CRC susceptibility.

**Graphical abstract:** 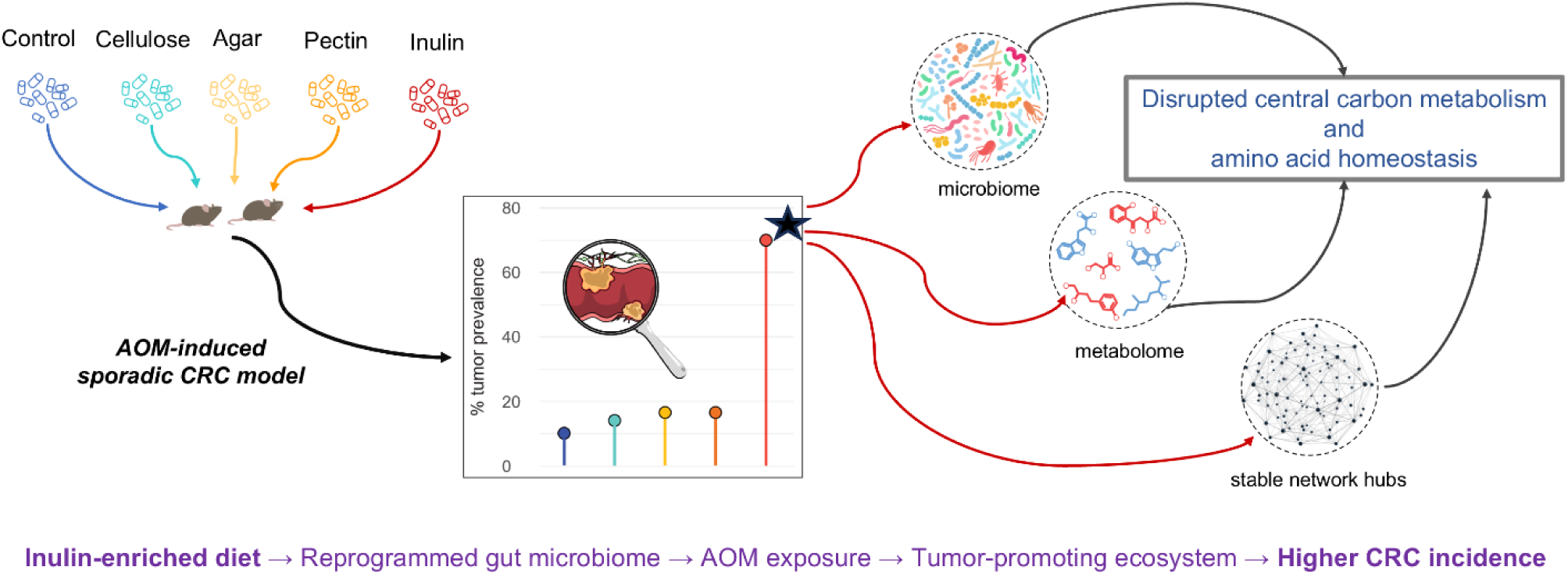

**Significance of this study:** What is already known on this subject?

- Dietary fiber from whole foods is consistently associated with a reduced risk of colorectal cancer (CRC).
- Inulin, a highly fermentable fructan used widely as both a prebiotic supplement and a food additive, selectively expands putatively beneficial gut taxa and is generally regarded as promoting gut health.
- Structurally distinct dietary fibers differentially shape the gut microbiome–metabolome axis.

What are the new findings?

- Refined inulin supplementation dramatically elevated colorectal tumor prevalence (∼70%) compared with all other fiber-supplemented and control groups (∼10–20%) under AOM-induced carcinogenic conditions.
- Longitudinal microbiome profiling identified 36 inulin-specific microbial signatures converging on two primary axes: depletion of SCFA-producing and cross-feeding taxa, and reprogramming of amino acid and nitrogen metabolism.
- Cecal metabolomics confirmed succinate, fumarate, and lactate accumulation alongside propionate and amino acid depletion, exclusively under AOM conditions, establishing carcinogen-dependent amplification of a diet-preconditioned metabolic vulnerability.
- Cecal content extracts from inulin-fed mice increased the proliferation of human HT-29 colorectal cancer cells compared with extracts from cellulose-fed mice, providing functional evidence that the inulin-conditioned luminal environment supports tumor growth.
- Stable microbiome-metabolome co-occurrence hubs were identified exclusively in AOM-treated animals, absent in carcinogen-naive counterparts, demonstrating that carcinogen exposure consolidates inulin-driven community restructuring into a structurally defined tumor-promoting ecosystem.

How might it impact on clinical practice in the foreseeable future?

- Under carcinogenic conditions, inulin-driven gut microbiome restructuring reveals a broader dysbiotic context of depleted protective fermentation capacity and oncometabolite accumulation, challenging its framing as a prebiotic benefit.
- Given the increasing use of isolated inulin in processed foods and as a dietary supplement, alongside the rising incidence of early-onset colorectal cancer, prospective human studies examining the association between inulin intake and colorectal cancer risk are warranted.
- Dietary fiber type specificity should be considered in clinical recommendations and supplement guidance for individuals at elevated colorectal cancer risk.

## 1. INTRODUCTION

Colorectal cancer (CRC) ranks as the third most commonly diagnosed malignancy and the second leading cause of cancer-related death worldwide, with global incidence projected to increase by approximately two-thirds by 2025 (1). The majority of CRC cases are sporadic, occurring without a defined hereditary syndrome or family history (2). Of particular concern is the sustained rise in early-onset CRC (EOCRC), diagnosed before age 50, whose incidence continues to increase even as rates among older adults have declined or stabilized, coinciding with expanded colorectal screening programs in Canada and the United States (3–5). A widely accepted explanatory framework implicates unfavorable environmental exposures beginning early in life, including consumption of processed foods (6,7).

Dietary fibers (DFs) are among the most extensively studied dietary factors associated with CRC risk. A dose-response meta-analysis reported an approximately 10% reduction in CRC risk for every 10 g/day increment in total dietary fiber intake (8). However, this protective effect is not uniform across fiber types; grain- and legume-derived fibers are consistently associated with lower CRC risk, whereas vegetable- and fruit-derived fiber consumption shows dose-dependent and divergent associations (9). This heterogeneity is attributed to structural properties including monosaccharide composition, glycosidic linkage chemistry, solubility, and viscosity, which collectively determine microbial accessibility and fermentation characteristics (10).

The gut microbiome represents the most direct interface between dietary fiber and host biology, fermenting fiber substrates into bioactive metabolites, modulating mucosal immune tone, and maintaining epithelial barrier integrity. Alterations in the gut microbiome are increasingly recognized as a hallmark of CRC, with distinct microbial profiles observed in patients with adenocarcinoma relative to healthy individuals and adenoma patients, implicating microbial ecological changes in the transition from adenoma to carcinoma (11–13). Collectively, these observations support a model in which the gut microbiome is fundamental to CRC-associated changes and the molecular environment that promotes tumorigenesis.

Despite growing recognition of the fiber-microbiota-host axis, how structurally distinct dietary fibers influence tumorigenic susceptibility, particularly when exposure begins early in life, remains poorly understood. Here, we used an azoxymethane (AOM)-induced mouse model of sporadic colorectal carcinogenesis to determine whether structurally distinct fibers; cellulose, agar, pectin, and inulin, differentially modulate colon tumor development. These DFs represent markedly different physicochemical and fermentative properties: cellulose is a linear, insoluble β-(1→4)-glucan; agar is a galactose-based gel-forming polysaccharide; pectin is a structurally heterogeneous polymer enriched in galacturonic acid; and inulin is a highly soluble and rapidly fermentable fructan. Nutritionally matched diets supplemented with this panel enabled evaluation of fiber-specific effects on tumorigenesis while minimizing confounding by caloric intake or macronutrient composition. Dietary interventions were initiated in 4-week-old mice, with carcinogen exposure beginning at 8 weeks, developmental stages corresponding approximately to 8 and 17 years of age in humans, respectively (14), providing an opportunity to examine whether early-life fiber exposure establishes divergent intestinal microbial states that subsequently modify carcinogenic susceptibility. We hypothesized that structurally distinct fibers would differentially shape the gut microbiome and its metabolic output, creating distinct colonic microenvironments associated with differential CRC susceptibility.

## 2. RESULTS

### 2.1. Structurally diverse dietary fibers exert distinct effects on colon tumor development in AOM-treated mice

We evaluated the effects of structurally distinct dietary fibers on sporadic colon cancer in an AOM-induced mouse model. The selected fibers span diversity in predominant backbone sugars and glycosidic linkage patterns, encompassing three structural polysaccharides (cellulose-CCD, agar-AgD, and pectin-PCD) and one storage polysaccharide (inulin-ICD). Mice were fed isocaloric, nutrient-matched diets supplemented with 7.5% w/w of each fiber or a low-fiber control diet for 28 weeks and received eight AOM injections (7.5 mg/kg, i.p.) at indicated timepoints (**Fig. 1A**). Throughout the study, mice in the pectin group maintained higher body weight than those in the inulin and control groups (**Fig. 1B**).

**Figure 1:**
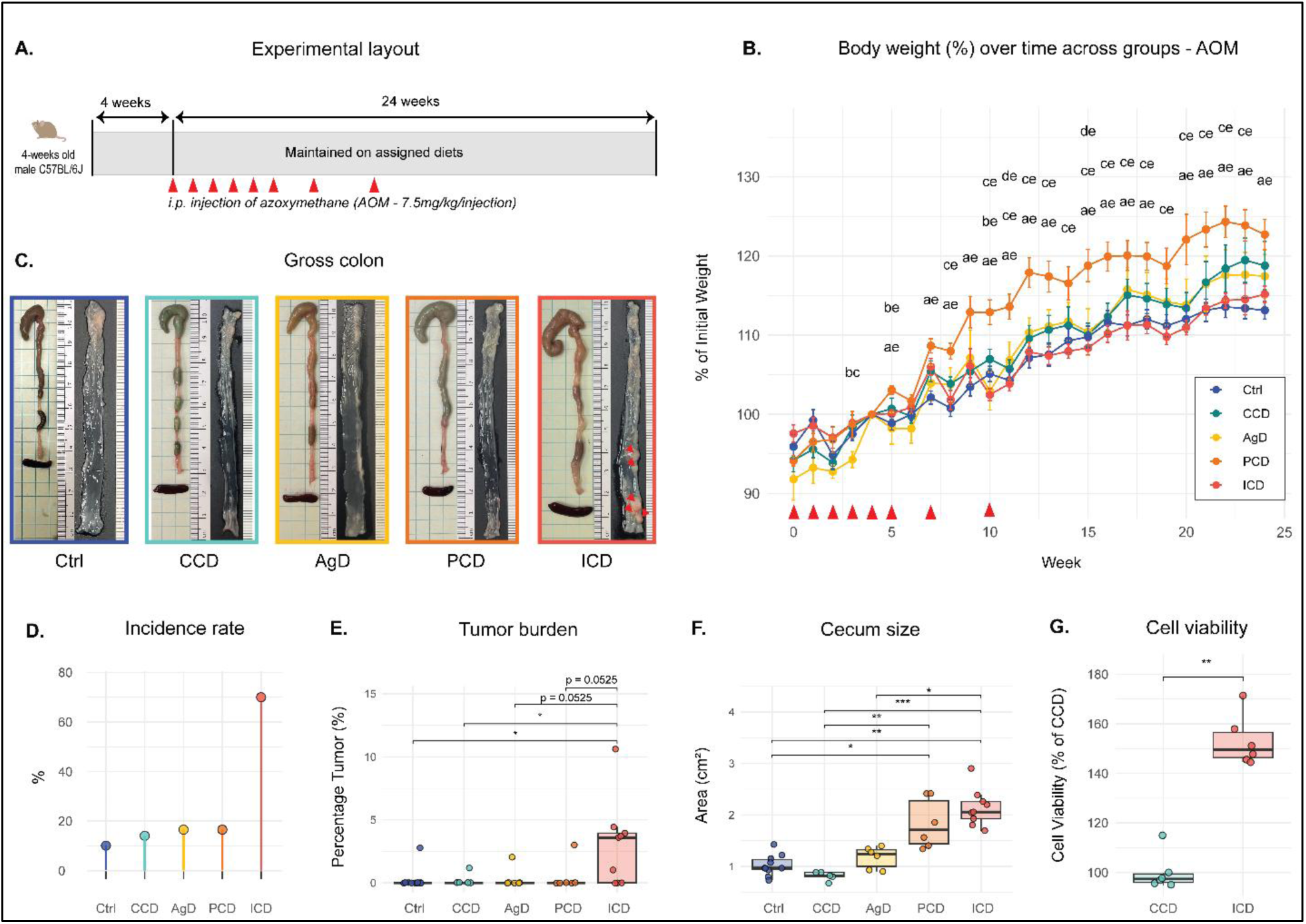
Phenotypic assessment of the AOM cohort. **A.** Study timeline. **B.** Percentage changes in body weight relative to week 4. **C.** Representative colon image. **D.** Tumor incidence. **E.** Tumor burden. **F.** Cecum size. **G.** Cell viability expressed as percentage relative to the CCD extract. Data are presented as individual datapoints with bars representing mean ± SEM. Statistical analyses were performed using non-parametric tests: Kruskal–Wallis tests followed by Dunn’s multiple-comparisons test with Benjamini–Hochberg (BH) correction for comparisons among more than two groups, and Wilcoxon rank-sum tests for comparisons between two groups. For body-weight measurements, pairwise significant comparisons at p-value 0.05 (BH) are denoted using symbols: Control (Ctrl) = a, Agar (AgD) = b, Cellulose (CCD) = d, Pectin (PCD) = e, and Inulin (ICD) = c.

Most notably, AOM-treated mice receiving the inulin-supplemented diet developed extensive colon tumorigenesis, evidenced by higher tumor incidence and larger tumor burden compared with other dietary groups (**Fig. 1C–E; Fig.-S1**). Inulin-induced exacerbation of sporadic colon tumorigenesis was reproduced in an independent cohort using the same experimental design, and data from both cohorts were pooled for subsequent analyses. In contrast, tumor incidence in mice fed cellulose-, agar-, or pectin-supplemented diets was comparable to the control group, indicating these fibers did not significantly influence colon tumorigenesis (**Fig. 1D–E**). Consistently, cecum size was similar in cellulose and control groups, with a trend toward enlargement in the agar group and significant cecal expansion in inulin- and pectin-supplemented groups, reflecting more active bacterial fermentation (**Fig. 1F**).

To determine whether luminal metabolites mediate the inulin-specific tumorigenesis effects, cecal content extracts from ICD and CCD mice, representing high- and low-incidence groups, respectively, were applied to the HT29 human colon cancer cell line. Approximately 50% more cells were observed in the ICD extract-treated group, indicating that ICD luminal metabolites promote cell proliferation relative to CCD (**Fig. 1G**). Histological analysis showed that colon tumors (inulin group) exhibited features of adenocarcinoma (**Fig. 2A**). Specifically, epithelial cells within tumor regions displayed enlarged, hyperchromatic nuclei with nuclear crowding and stratification. Acidic mucin staining was comparable in normal colonic tissue across groups, whereas tumor regions showed a significant reduction in acidic mucin staining (**Fig. 2B**). Importantly, no macroscopically visible colon tumors were detected in any fiber-only group in the absence of AOM (**Fig. S2**), indicating that dietary fiber increased susceptibility to colon cancer only in the presence of a secondary carcinogenic trigger.

**Figure 2:**
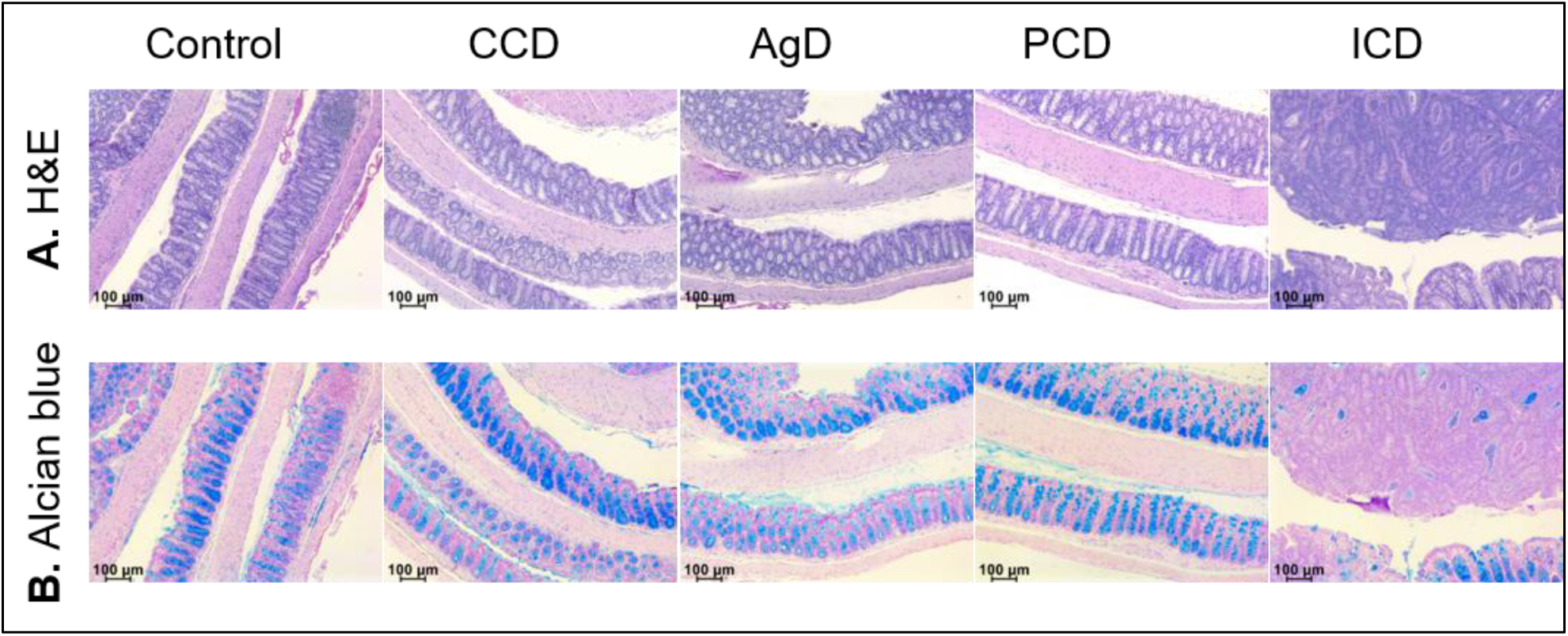
Colon tumors in the inulin-supplemented group exhibited features of adenocarcinoma, and loss of acidic mucin. Representative H&E (upper panel) and Alcian blue (lower panel)–stained images from AOM treated control and DF intervention groups. Tumor regions in inulin group showed loss of crypt architecture, irregular and branched glands, goblet cell depletion, and enlarged, pleomorphic nuclei. Acidic mucin was markedly reduced in tumor regions.

### 2.2. Diet-specific temporal microbiome dynamics identify inulin-associated microbial signatures

To establish whether the inulin-specific tumor phenotype was underpinned by distinct gut microbiome dynamics, we performed longitudinal 16S rRNA gene profiling of fecal samples collected across seven timepoints throughout the AOM carcinogenesis window. NMDS ordination of Bray-Curtis dissimilarity matrices revealed community-level separation associated with both dietary group and timepoint (Fig. S3). PERMANOVA confirmed significant marginal contributions of diet (R² = 0.237, p = 0.001), timepoint (R² = 0.130, p = 0.001), and their interaction (R² = 0.104, p = 0.001) to overall community variation, indicating that dietary fiber composition drives diverging temporal microbiome trajectories rather than static compositional differences. Cancer incidence group was independently associated with community composition (R² = 0.123, p = 0.001).

Linear mixed model analysis identified 72 genera with significant diet × time interactions (FDR adjusted p-value < 0.05; **Table-S1**). ICD-specific signatures were defined as taxa showing persistent beta-coefficient divergence from the combined low-incidence fiber group (AgD, CCD, PCD) at four or more of seven sampling timepoints, with ICD exhibiting a uniquely extreme predicted abundance at one or more timepoints, yielding 36 ICD-specific microbial signatures (**Fig. 3**; individual taxon plots in **Figs. S4–S6**). The majority of ICD-specific taxa maintained consistent directional shifts, either persistently higher or lower than other fiber groups, across four or more of seven sampling timepoints, reflecting stable diet-driven compositional states. Fifteen taxa showed consistently lower predicted abundance in ICD relative to other fiber groups, including *Alistipes*, *Butyricicoccus*, *Acetatifactor*, *Acetitomaculum*, *Eubacterium*, *Clostridium_Q*, and Lachnospiraceae_1XD42-69, among others (**Fig. 3**). These taxa encompass phylogenetically diverse butyrate and acetate producers, suggesting a broad and coordinated reduction in SCFA-producing capacity (15–20). *Emergencia* and Eggerthellaceae_QWKK01 were also consistently depleted, representing taxa associated with diverse anaerobic metabolic processes including fermentation and amino acid metabolism (21,22).

**Figure 3:**
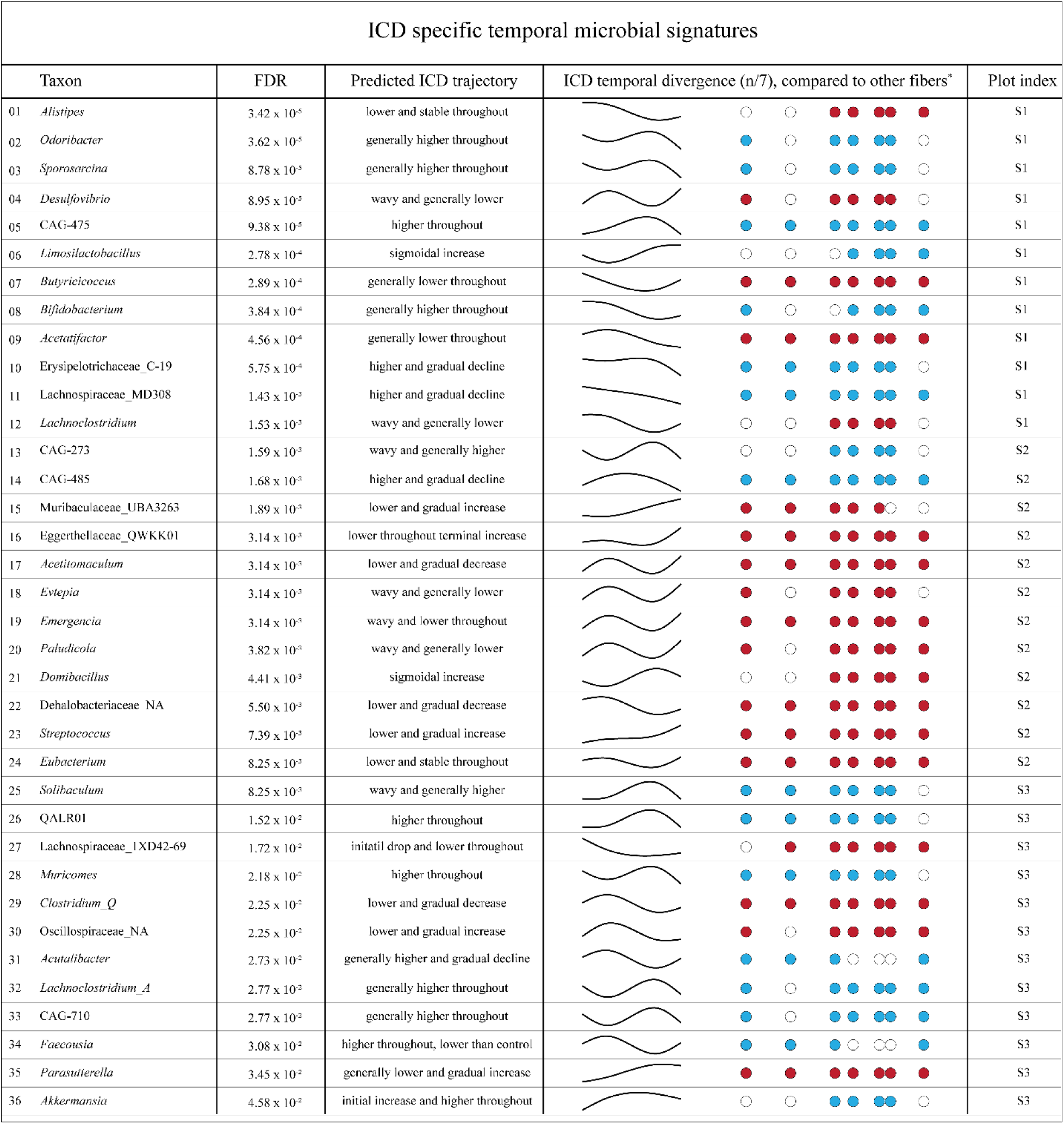
ICD-specific temporal microbial signatures identified by linear mixed model analysis. Thirty-six genera exhibiting significant ICD-specific temporal divergence from the combined low-incidence fiber group (AgD, CCD, PCD) were identified from 72 genera with significant diet × time interactions (p-value < 0.05) in a linear mixed model with a natural cubic spline for time, as described in methods, applied to longitudinal 16S rRNA gene profiling data from AOM-treated mice across seven timepoints (weeks 0–24). Taxa are ranked by p-value and annotated with their model-estimated CLR abundance trajectory for ICD, derived from the fitted linear mixed model. ICD temporal divergence at each of seven sampling timepoints (n = W0, W06, W12, W14, W18, W19, W24) is depicted as a dot plot, where filled blue circles indicate timepoints at which ICD abundance was significantly higher than the mean of other fiber groups, filled red circles indicate significantly lower abundance, and open circles indicate non-significant divergence. Trajectory sketches provide an abstract representation of the ICD beta-coefficient trend across the experimental timeline. Individual predicted trajectory and beta-coefficient plots for these listed taxa are provided in the supplementary data files (**Fig. S4-S6**).

Twenty-one taxa were enriched in ICD relative to other fiber groups, exhibiting sustained elevation, gradual increase, or sigmoidal dynamics (**Fig. 3**). Among the most significantly enriched were CAG-475, showing consistently higher abundance across all seven timepoints, *Bifidobacterium*, *Sporosarcina*, and *Akkermansia*. Additional enriched taxa included multiple Muribaculaceae-affiliated genera, *Lachnoclostridium*, *Muricomes*, and *Solibaculum*. Several taxa including Erysipelotrichaceae_C-19, Lachnospiraceae_MD308, and CAG-485 showed higher initial abundance followed by gradual decline, suggesting early ICD-driven expansion that subsequently moderated as the experiment progressed.

Temporal divergence was most pronounced at mid-to-late timepoints (weeks 12–24), consistent with progressive dietary shaping of the microbiome across the carcinogenesis window. The breadth and temporal consistency of ICD-specific signatures across functionally diverse microbial lineages, spanning fermentation, cross-feeding, amino acid metabolism, and immune-associated taxa, indicates a coordinated community-level reorganization. These specific microbial signatures displayed broader biological links associated with tumorigenesis, suggesting ICD-induced microbial modulation as a potential factor underlying the higher tumor prevalence observed for this diet.

### 2.3. Cecal metabolomics identifies inulin- and AOM-associated disruption of fermentation outputs and amino acid pools

To assess the functional metabolic consequences of diet-driven microbiome reorganization at the terminal timepoint, cecal metabolite concentrations were profiled by ¹H NMR spectroscopy across AOM-treated and BASAL cohorts at terminal timepoint. Principal coordinates analysis of the cecal metabolome revealed partial separation between dietary groups and between treatment conditions (**Fig. S7)**, with PERMANOVA confirming significant contributions of both diet and treatment to overall metabolome composition (p-value < 0.05). Among 38 quantified metabolites, 18 showed significant differential abundance (p-value < 0.05) in at least one ICD-involving comparison within the AOM cohort, with no metabolites reaching significance in any BASAL comparison (**Fig. 4**; **Table-S2**). This treatment-dependent alteration, visible across metabolite panels (**Fig. 4)**, indicates that the observed metabolic shifts reflect diet-preconditioned states that are substantially consolidated by carcinogen exposure rather than arising *de novo* as a consequence of tumor development.

**Figure 4:**
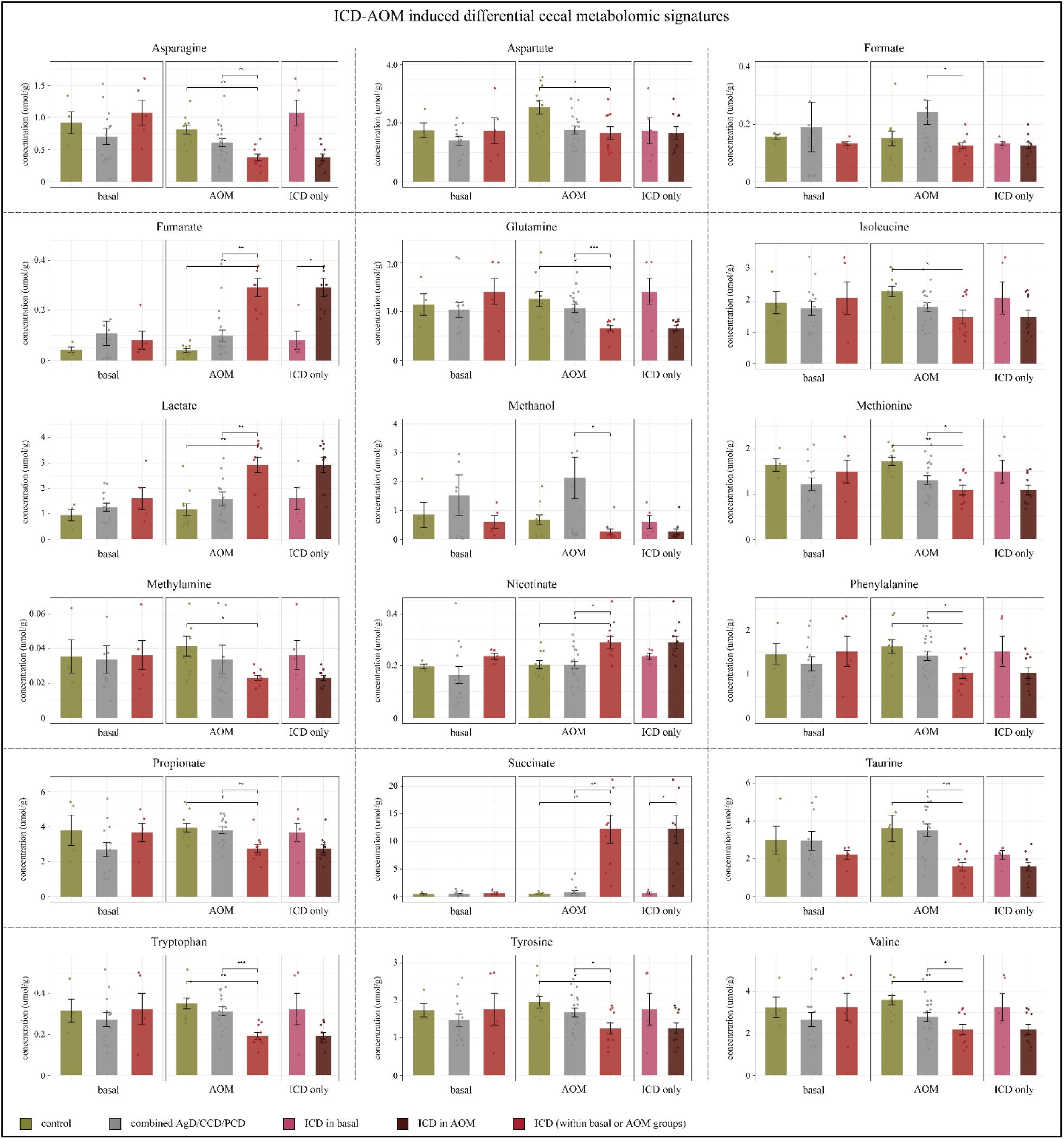
ICD- and AOM-associated differential cecal metabolite concentrations at the terminal timepoint. Cecal concentrations (µmol/g) of 18 metabolites showing significant differential abundance (p-value < 0.05) in at least one ICD-involving comparison, are presented as bar plots (mean ± SE) with individual data points overlaid. Each metabolite panel comprises three comparison groups: BASAL (left), AOM (center), and ICD specific (right). Within BASAL and AOM panels, bars represent the unamended Control (olive), combined low-incidence fiber group AgD/CCD/PCD (grey), and ICD (red). The ICD-only panel compares ICD-BASAL (pink) and ICD-AOM (brown) directly. Significance brackets indicate pairwise comparisons passing p-value < 0.05 (\**), p-value < 0.01 (**)**, or p-value < 0.001 (\*\*\****) by Welch’s two-sample t-test with Benjamini-Hochberg correction. Brackets are shown only for significant comparisons. Individual data points are colored red for ICD samples and shown in grey for all other groups. Full differential abundance results for all 38 quantified metabolites are provided in **Table-S2**.

The most pronounced differential signal was the significant elevation of succinate in ICD mice within the AOM cohort relative to both the unamended Control and the combined low-incidence fiber group (*AgD/CCD/PCD; p-value < 0.01 for both comparisons*). Fumarate was similarly elevated in ICD-AOM mice relative to both comparison groups (p-value < 0.01), with neither metabolite showing significant differences in the BASAL cohort, confirming that intermediate metabolite accumulation is amplified by carcinogen exposure and not solely attributable to inulin fermentation. Lactate was significantly higher in ICD relative to other fiber groups within AOM (p-value < 0.01) and trended higher in the ICD-only panel, consistent with increased anaerobic fermentation flux (23). Together, succinate, fumarate, and lactate represent the most statistically robust and biologically coherent cluster of metabolic differences associated with the ICD-AOM condition. In contrast, propionate was significantly reduced in ICD mice within the AOM cohort relative to the combined fiber group (p-value < 0.01), providing direct endpoint confirmation of reduced terminal fermentation output. Formate showed a significant reduction in ICD-AOM relative to other fibers (p-value < 0.05), with the ICD-only panel suggesting this reduction is present across both treatment conditions. Methanol was also significantly reduced in ICD-AOM mice relative to the combined fiber group (p-value < 0.05).

A second major category of differentially abundant metabolites comprised amino acids, with widespread depletion observed across multiple structural classes in ICD-AOM mice (**Fig. 4**). Tryptophan was among the most significantly depleted (p-value < 0.001 vs combined fibers within AOM), followed by glutamine (p-value < 0.001), taurine (p-value < 0.001), methionine (p-value < 0.05), phenylalanine (p-value < 0.05), isoleucine (p-value < 0.05), and valine (p-value < 0.05). Asparagine and aspartate showed significant reductions in ICD relative to other fiber groups within AOM (p-value < 0.01 and p-value < 0.05 respectively). Across this amino acid panel, the pattern was also consistent, differences were attenuated in the BASAL cohort and amplified under AOM conditions, reinforcing the diet-preconditioning interpretation. In addition, Nicotinate was significantly elevated in ICD-AOM mice relative to other fiber groups (p-value < 0.05), while methylamine was significantly reduced (p-value < 0.05), consistent with altered nitrogen metabolism dynamics.

Collectively, the cecal metabolome at week 24 identified two functionally coherent categories of ICD-associated metabolic disruption, accumulation of central carbon intermediates and depletion of amino acid pools, both of which were substantially amplified under AOM treatment. These observations provide endpoint metabolite-level corroboration of the temporal microbiome signatures described above, and their mechanistic integration is addressed in the Discussion following multi-omics network analysis. Importantly, the complete absence of significant metabolite differences in the BASAL cohort, despite identical dietary interventions and equivalent statistical power, indicates that the ICD-associated cecal metabolic disruptions identified here are not a direct consequence of fiber composition alone, but emerge specifically under AOM-induced carcinogenic pressure. This observation is consistent with a model in which ICD preconditioning of the gut microbiome establishes a latent metabolic vulnerability that is converted into measurable biochemical disruption upon carcinogen exposure, a model whose structural basis is resolved in the microbiome-metabolome network analysis described below.

### 2.4. Multi-omics network integration identifies stable microbiome-metabolome co-occurrence modules exclusive to AOM-treated animals

To determine whether the microbiome and metabolomics signatures described above reflect coordinated biological interactions, Spearman rank correlations were computed between CLR-transformed genus-level microbiome abundances and CLR + Pareto-scaled cecal NMR metabolite concentrations at week 24, independently for AOM-treated (*n = 39*) and BASAL (*n = 22*) cohorts. Co-occurrence networks were constructed from significant correlations (*|ρ| > 0.449, FDR p-value < 0.1 for AOM; |ρ| > 0.611, FDR p-value < 0.1 for BASAL*), yielding 251 edges across 13 clusters for AOM and 145 edges across 5 clusters for BASAL (**Fig. 5A-B**). Community detection by ClusterOne identified six clusters in the AOM network and three in the BASAL network meeting the significance threshold (*p-value < 0.05*) (24). Cluster robustness was evaluated by Jaccard similarity-based bootstrap stability analysis across three progressively relaxed correlation thresholds, classifying hubs as Strong, Moderate, or Weak based on membership persistence across tiers (25). Two Strong hubs, clusters whose membership remained stable across all three bootstrap tiers (Jaccard ≥ 0.3), were identified exclusively in the AOM cohort (**Fig. 5C**). No Strong hubs were detected in the BASAL cohort, where all clusters showed low Jaccard stability and were classified as Weak or Unstable. This asymmetry indicates that stable, reproducible microbiome-metabolome co-regulatory modules emerge specifically under AOM-induced carcinogenic conditions and are not a direct consequence of dietary fiber composition alone.

**Figure 5:**
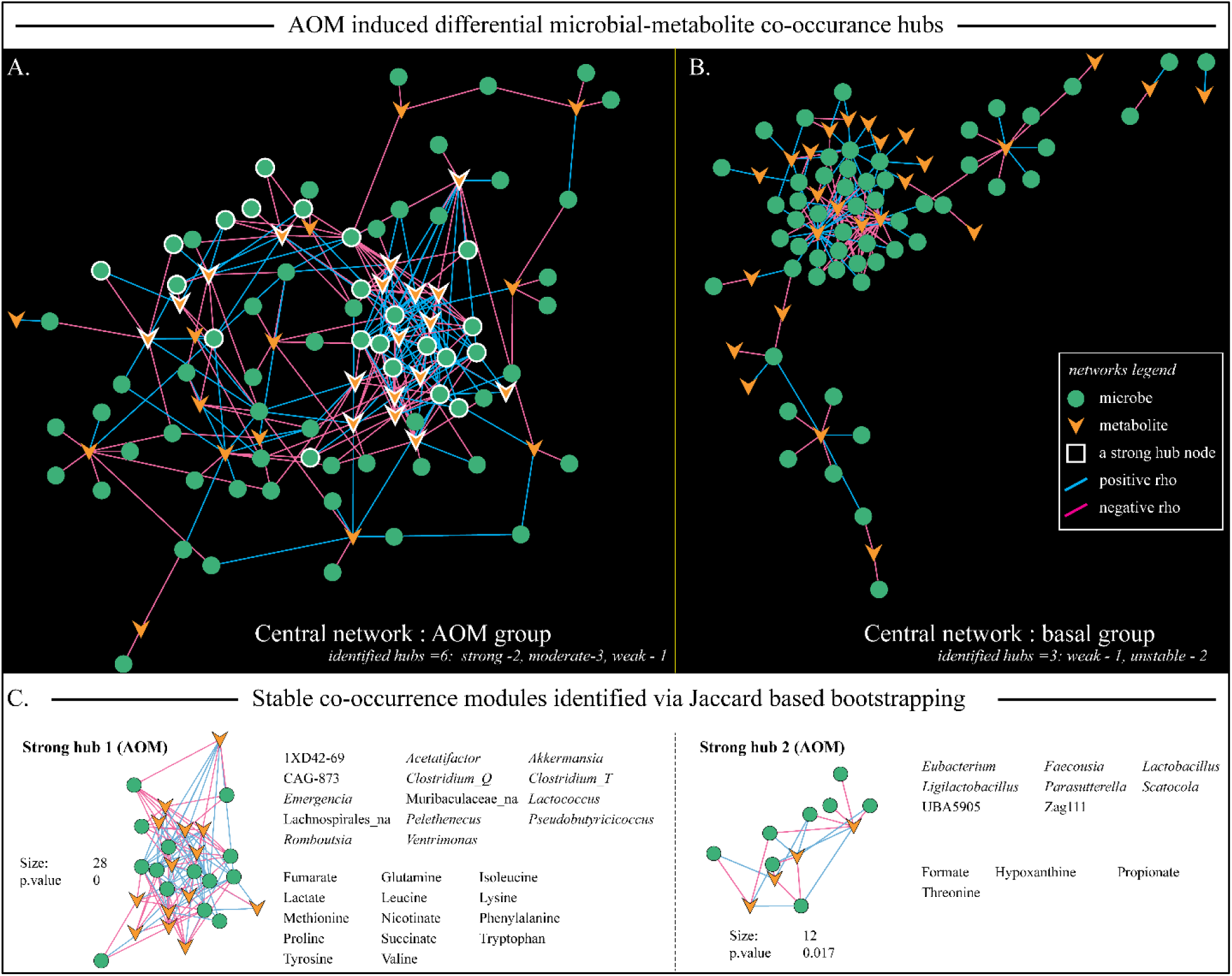
AOM-induced stable microbiome-metabolome co-occurrence hubs identified by network integration and Jaccard-based bootstrap stability analysis. (A) Co-occurrence network for AOM-treated animals (n = 39) constructed from significant Spearman rank correlations between CLR-transformed genus-level microbiome abundances and CLR + Pareto-scaled cecal NMR metabolite concentrations at week 24 (|ρ| > 0.449, p-value < 0.1). Nodes represent individual microbial genera (green circles) or metabolites (orange triangles). Edges represent significant FDR-adjusted pairwise correlations, colored blue for positive associations and pink for negative associations. Six clusters were identified by ClusterOne community detection; two Strong hubs identified by Jaccard-based bootstrap stability analysis are indicated by white-outlined nodes. (B) Corresponding co-occurrence network for carcinogen-naive BASAL animals (n = 22; |ρ| > 0.611, p-value < 0.1). Three clusters were identified, none of which met the stability threshold for Strong hub classification across bootstrap tiers. (C) Detailed membership of the two AOM Strong hubs identified by Jaccard similarity-based bootstrap stability analysis across three progressively relaxed correlation thresholds (AOM tiers: |ρ| > 0.40, 0.35, 0.30 without p-value correction). Strong Hub 1 (n = 28 members, p ≈ 0) and Strong Hub 2 (n = 12 members, p = 0.017) are shown as subnetwork visualizations with full member lists. Hub size and ClusterOne permutation p-value are indicated below each subnetwork.z

AOM Strong Hub 1 was the largest and most significant cluster identified (*n = 28 members, p ≈ 0;* **Fig. 5C**). Microbiome members included *Akkermansia*, *Pseudobutyricicoccus*, *Ventrimonas*, *Emergencia*, *Romboutsia*, *Pelethenecus*, *Lactococcus*, *Clostridium_T*, *Acetatifactor*, *Clostridium_Q*, Muribaculaceae_NA, 1XD42-69, and CAG-873, among others. Metabolite members comprised succinate, fumarate, lactate, and a broad panel of amino acids including tryptophan, tyrosine, phenylalanine, methionine, lysine, leucine, isoleucine, valine, glutamine, proline, and nicotinate. The co-occurrence of fermentation intermediates and diverse amino acids within a single stable interaction module indicates structural coupling between central carbon disruption and amino acid pool depletion at the network level. AOM Strong Hub 2 comprised 12 members (*p = 0.017;* **Fig. 5C**), with microbiome members including *Eubacterium*, *Faecousia*, *Parasutterella*, *Ligilactobacillus*, *Scatocola*, UBA5905, and Zag111, and metabolite members comprising formate, propionate, hypoxanthine, and threonine. The co-occurrence of propionate and formate, both reduced in ICD-AOM mice in the differential metabolomics analysis, with fermentation-associated taxa within a stable module identifies a second structurally distinct but complementary interaction network centered on disrupted terminal fermentation outputs. Three Moderate hubs were also identified in the AOM network, stable across two of three bootstrap tiers (**Table-S3**), with biological content spanning immune-associated taxa, purine metabolism markers, and sulfur and one-carbon metabolism members. No interpretable stable structure was recovered from the BASAL network beyond the Weak and Unstable classifications noted above.

Notably, several microbiome members of both AOM Strong hubs, including *Akkermansia*, *Emergencia*, *Acetatifactor*, *Clostridium_Q*, *Eubacterium*, 1XD42-69, *Pseudobutyricicoccus*, and *Ventrimonas*, were independently identified as ICD-specific temporal signatures in the longitudinal microbiome analysis (**Fig. 3**). Similarly, metabolite members of Hub 1, including succinate, fumarate, lactate, and multiple amino acids, corresponded directly to the differentially abundant metabolites identified in the AOM cecal metabolomics analysis (**Fig. 4**). This structural overlap between the AOM-wide network architecture and the ICD-specific signatures identified across both preceding analytical layers provides convergent multi-omics evidence for the two primary biological axes addressed in the Discussion.

## 3. DISCUSSION

Processed DF supplementation has emerged as an attractive strategy for increasing daily fiber intake. However, such DF supplementation are typically consumed as a single, purified type that is highly accessible to gut microbial metabolism, which can drive a rapid and sometimes unfavorable shift in gut microbiome composition and metabolic output. Consistent with this, a meta-analysis of 25 prospective studies found that CRC risk reduction is observed primarily with fiber-containing foods, not with fiber intake as an isolated nutrient (26). In whole or minimally processed foods, soluble dietary fibers are largely contained within intact plant cell wall matrix that meters their release to the microbiota; refined mono-fiber supplementation removes that metering step entirely, delivering a concentrated, immediately fermentable substrate bolus with no dietary precedent. Our findings demonstrated that supplementation with inulin, a highly fermentable DF, raised colon tumor incidence to ∼70%, compared with ∼20% for cellulose, agar, or pectin, and 10% for control. Since both agar and pectin are also soluble and fermentable, the higher tumor incidence in the inulin group implicates the specific fermentation trajectory imposed by a β-(2→1)-linked fructan storage polysaccharide as a primary contributor to colon tumor development. The inulin-specific microbiome signatures resolved onto two axes: loss of SCFA-producing and cross-feeding taxa, and reprogramming of amino acid and nitrogen metabolism; together describing a community unable to complete saccharolytic fermentation and defaulting toward proteolytic routes. Cecal metabolomic profiling corroborated these findings, revealing luminal accumulation of succinate, fumarate, and lactate alongside a reduction in propionate. These changes indicate a breakdown of metabolic cross-feeding networks rather than excessive fermentation, given that succinate is typically rapidly metabolized to propionate within the gut lumen (27). The emergence of this metabolic disruption exclusively under AOM treatment suggests that inulin primes the intestinal ecosystem for metabolic instability that becomes manifest as tumorigenesis following carcinogen challenge.

### 3.1. Inulin-induced preconditioning of the gut microbiome as a determinant of AOM-induced colorectal cancer prevalence

Among the four dietary conditions tested, the inulin-supplemented diet was associated with markedly higher colorectal tumor prevalence under AOM-induced carcinogenesis, in contrast to the low and comparable tumor rates observed across all other fiber-supplemented diets and the unamended control. This differential phenotypic outcome, in which three structurally and compositionally distinct dietary interventions produced comparable cancer prevalence while one diverged substantially, indicates that the elevated tumor burden in ICD-fed animals reflects a diet-specific biological effect rather than a general consequence of fiber supplementation or AOM exposure. Given that all groups received identical carcinogen doses under equivalent housing conditions, the source of this divergence was investigated through the gut microbiome as the primary diet-responsive biological system.

To interrogate the mechanistic basis of ICD-associated tumor prevalence, three complementary analytical layers were employed, each addressing a distinct dimension of the diet-microbiome-cancer relationship. Longitudinal 16S rRNA gene profiling of AOM-treated animals across seven timepoints captured the temporal dynamics of microbiome reorganization under sustained dietary pressure, identifying ICD-specific compositional trajectories that diverge progressively from low-incidence fiber groups across the carcinogenesis window. Endpoint cecal metabolomics independently assessed the functional metabolic consequences of these community shifts at the terminal timepoint in both AOM-treated and carcinogen-naive BASAL animals, enabling discrimination between diet-preconditioned metabolic states and those arising specifically under carcinogenic conditions. Microbiome-metabolome network integration then resolved the structural co-regulatory architecture of these interactions at the endpoint, providing a systems-level view of how diet-driven microbial shifts translate into coordinated metabolic reorganization. Together, these three layers address temporally distinct aspects of the same biological process, community establishment, functional output, and interaction architecture, and their convergence on consistent biological signals substantially strengthens the mechanistic inferences drawn from any single analytical approach.

Across all these layers, the data converges on two primary biological axes that are coherently and independently supported by microbiome temporal dynamics, endpoint metabolomic shifts, and stable network co-occurrence modules. A third set of observations, encompassing immune modulation, ecological restructuring driven by inulin fermentation, and sulfur metabolism redistribution, emerged as secondary but internally consistent signals whose contribution to ICD-associated carcinogenesis warrants further investigation. Collectively, these findings support a model in which the inulin content of the ICD diet preconditions the gut microbiome toward a state of functional vulnerability, characterized by disrupted fermentation throughput and reprogrammed amino acid metabolism, that is subsequently amplified and structurally consolidated by AOM-induced carcinogenic pressure into the tumor-promoting ecosystem reflected in the higher cancer prevalence observed phenotypically. The mechanistic basis of each axis, and its convergent support across analytical layers, is addressed in the following sections.

### 3.2. Axis 1: Disruption of fermentation throughput and accumulation of central carbon intermediates

The most consistently supported biological axis across all three analytical layers was a coordinated disruption of microbial fermentation throughput, characterized by concurrent depletion of terminal fermentation capacity and accumulation of central carbon intermediates. These two features are mechanistically linked as upstream and downstream consequences of the same ecological disruption: the loss of cross-feeding consumers that normally convert fermentation intermediates into SCFAs (27).

In the longitudinal microbiome analysis, multiple phylogenetically diverse butyrate and acetate producers were consistently depleted in ICD mice relative to the combined low-incidence fiber group across the majority of sampling timepoints, including *Butyricicoccus*, *Acetatifactor*, *Acetitomaculum*, *Eubacterium*, *Clostridium*, Lachnospiraceae_1XD42-69, and multiple Oscillospiraceae-affiliated genera (15–20). Their collective depletion indicates a broad reduction in SCFA-producing capacity rather than competitive exclusion of individual lineages. Crucially, the loss of *Emergencia*, a taxon identified as a central node in the AOM multi-omics network and linked to fermentation cross-feeding through its connectivity to succinate, lactate, and amino acid metabolism, likely represents the disruption of a metabolic coordinator that would normally facilitate conversion of fermentation intermediates into terminal products, creating conditions for intermediate accumulation independently of any change in production rates (28,29). *Akkermansia*, by contrast, was consistently enriched across multiple timepoints. Its enrichment is most appropriately interpreted as a marker of increased host-derived substrate utilization under conditions of depleted saccharolytic fermentation, with succinate generated through mucin catabolism contributing secondarily to the accumulating intermediate pool (30,31).

The endpoint cecal metabolomics data provides direct biochemical confirmation of this model. Succinate and fumarate were significantly elevated in ICD mice within the AOM cohort relative to both the unamended control and the combined low-incidence fiber group, whereas neither metabolite differed significantly among BASAL animals. Thus, in this experimental context, the intermediate accumulation requires AOM-induced carcinogenic pressure to become biochemically manifest rather than arising passively from inulin fermentation, consistent with our prior observations (32). Lactate was similarly elevated under AOM conditions, consistent with increased anaerobic fermentation flux, while propionate was significantly reduced, providing direct endpoint corroboration of depleted SCFA-producing capacity (27). The absence of significant butyrate and acetate differences is most parsimoniously explained by altered host consumption dynamics: colonocyte butyrate oxidation is progressively downregulated in colorectal tumor cells adopting glycolytic metabolism consistent with the Warburg effect, allowing cecal concentrations to remain stable despite reduced microbial production (33,34).

At the network level, Strong Hub1 contains succinate, fumarate, lactate, and several ICD-depleted cross-feeding taxa, including *Emergencia*, *Acetatifactor*, *Clostridium_Q*, and 1XD42-69, within a single stable co-occurrence module, confirming that their co-variation is a reproducible structural feature of the AOM-associated gut ecosystem. Strong Hub2 independently captures propionate and formate alongside fermentation-associated taxa including *Eubacterium*, *Parasutterella*, and *Faecousia*, representing a complementary module centered on disrupted terminal fermentation outputs. The structural separation of these two hubs; one organized around intermediate accumulation, the other around terminal product depletion, reflects the mechanistic layering of the same underlying disruption. Critically, neither hub was detected in the BASAL network, confirming these co-regulatory structures are carcinogen-dependent manifestations of the diet-preconditioned state.

The biological significance of succinate accumulation extends well beyond its role as a fermentation byproduct. Extracellular succinate activates SUCNR1 on immune and epithelial cells to promote pro-inflammatory NF-κB signaling, while elevated intracellular succinate inhibits prolyl hydroxylase enzymes responsible for HIF-1α degradation, stabilizing downstream targets including VEGF, glucose transporters, and glycolytic enzymes that directly support tumor angiogenesis and metabolic reprogramming (35,36). The microbiome-driven accumulation of succinate in the ICD cecal environment therefore represents a mechanistically plausible route through which diet-driven ecological disruption translates into a tumor-promoting biochemical milieu.

### 3.3. Axis 2: Reprogramming of amino acid and nitrogen metabolism

The second consistently supported biological axis was a coordinated reprogramming of amino acid and nitrogen metabolism, characterized by depletion of taxa involved in amino acid fermentation and nitrogen coupling, widespread reduction of cecal amino acid concentrations, and structural integration of amino acid pools within the primary AOM-associated network hub. This axis emerged as mechanistically coupled to Axis 1, the same ecological disruption that depletes fermentation cross-feeders also dismantles the microbial machinery responsible for balanced amino acid utilization, and the network analysis confirms that amino acid depletion and central carbon intermediate accumulation are structurally co-organized within a single stable interaction module.

In the longitudinal microbiome analysis, an uncultivated genus within the Eggerthellaceae family (designated as QWKK01) was among the most significantly depleted taxa. Members of the Eggerthellaceae family are structurally recognized for their metabolic versatility in the healthy gut, and their depletion is a frequent hallmark of the shift from cancer-free to CRC cohorts (37). *Emergencia*’s depletion, discussed in Axis 1 as a fermentation cross-feeder, additionally represents a dual disruption spanning nitrogen flux. *Eubacterium*, similarly depleted across most timepoints, has established roles in amino acid fermentation and nitrogenous substrate integration into SCFA synthesis pathways (16,27). In contrast, *Solibaculum* was enriched and, had been associated with taurine and hypotaurine metabolism, thus suggesting a shift in sulfur-containing amino acid routing toward alternative metabolic destinations (38). *Desulfovibrio*, canonically elevated in CRC cohorts (39), was paradoxically depleted, most likely explained by substrate competition from high inulin fermentability rather than a tumor-associated mechanism, illustrating the importance of dietary context in shaping microbiome-disease associations (40).

The cecal metabolomics data provides direct validation of microbiome-predicted amino acid flux disruption. Widespread depletion was observed across aromatic amino acids (tryptophan, tyrosine, phenylalanine), branched-chain amino acids (valine, isoleucine), nitrogen-carrying amino acids (asparagine, glutamine), and sulfur-containing methionine. Taurine, with established roles in bile acid conjugation and mucosal cytoprotection (41), was among the most significantly depleted (p-value < 0.001). Its reduction shifts bile acid conjugation toward glycine-conjugated species with altered antimicrobial and receptor-signaling properties, compounding epithelial vulnerability through a mechanism independent of the fermentation disruptions described in Axis 1 (41). Critically, amino acid depletions were attenuated in the BASAL cohort and amplified under AOM conditions, mirroring the treatment-dependent pattern of Axis 1 and confirming the preconditioning model.

The mechanistic basis for widespread amino acid depletion is multifactorial. Reduced abundance of amino acid-fermenting taxa diminishes microbial contributions to the cecal pool, while elevated succinate and fumarate concentrations are consistent with increased anaplerotic channeling of amino acid carbon skeletons, particularly from glutamine, aspartate, and branched-chain amino acids, into TCA cycle intermediates, creating a metabolic sink that mutually amplifies both axes (42,43). Glutamine depletion is of particular note, as it serves as the primary nitrogen donor for nucleotide biosynthesis and preferred fuel for rapidly proliferating cells, and its reduction may reflect both reduced microbial production and increased tumor cell demand (42). At the network level, the co-occurrence of succinate, fumarate, lactate, and a broad amino acid panel within AOM Strong Hub1, alongside ICD-depleted cross-feeding and amino acid-fermenting taxa, establishes that these disruptions are components of a structurally integrated reorganization rather than parallel independent signals. The complete absence of this hub from the BASAL network confirms its carcinogenic environment-dependent nature.

### 3.4. Coordinated broader signatures of microbiome restructuring

The two primary axes were accompanied by broader coordinated signatures that reinforce and contextualize the core metabolic reorganization. Inulin-driven *Bifidobacterium* expansion may act as the upstream ecological organizer of both axes, altering central fermentation outputs, including succinate, through bifidobacterial carbohydrate metabolism, while competitively displacing canonical SCFA producers, directly linking ICD fiber composition to the central carbon disruption of Axis 1 through a mechanistically traceable pathway (44). *Akkermansia* enrichment reflects a community-level shift toward host mucin utilization under conditions of depleted saccharolytic fermentation, potentially compounding epithelial vulnerability through mucus layer degradation (30,31). Its co-occurrence with succinate, fumarate, and ICD-depleted cross-feeders within AOM Strong Hub 1 establishes this shift as structurally integrated within the core dysbiotic module rather than a peripheral observation. Immune-modulatory taxa including CAG-475, which showed significant ICD-specific divergence at all seven sampling timepoints, and the CRC-associated biomarker *Lachnoclostridium* (45), collectively support a chronically altered mucosal immune environment whose pro-inflammatory tone is mechanistically downstream of the SCFA depletion and succinate accumulation described in Axes 1 and 2 (35).0

## 4. CONCLUSION

Inulin supplementation raised colorectal tumor prevalence to ∼70% under AOM-induced carcinogenesis, compared with ∼10–20% in all other fiber-supplemented and control groups. Convergent evidence across longitudinal microbiome profiling, endpoint cecal metabolomics, and microbiome-metabolome network integration identified two mechanistically coupled biological axes underlying this divergence: disruption of fermentation throughput with oncometabolite accumulation, and reprogramming of amino acid and nitrogen metabolism. Together these axes describe a gut ecosystem that is metabolically impoverished in its protective functions and enriched in pro-tumorigenic intermediates. The functional relevance of this luminal environment was directly demonstrated by the increased proliferation of HT29 colorectal cancer cells exposed to ICD cecal extracts relative to CCD, providing experimental support for the tumor-promoting capacity of the inulin-conditioned gut milieu. Critically, these metabolic and network-level signatures were absent in carcinogen-naive animals receiving identical diets, demonstrating that inulin fermentation creates a latent biological vulnerability that carcinogen exposure amplifies into a structurally consolidated, tumor-promoting ecosystem. This interpretation is structurally reinforced by the complete absence of stable microbiome-metabolome co-occurrence hubs in carcinogen-naive BASAL animals, demonstrating that inulin-driven community shifts alone are insufficient to generate the consolidated interaction architecture observed under AOM, and that carcinogen exposure is the consolidating agent that converts diet-preconditioned vulnerability into a structurally defined, tumor-promoting ecosystem. The higher tumor prevalence in ICD-fed animals therefore likely reflects not a greater carcinogenic insult but a more permissive biological context, one in which the same carcinogenic stimulus produces a more severe oncological outcome because the gut ecosystem has been preconditioned to offer less resistance.

Among the most mechanistically significant findings is the accumulation of succinate in the ICD cecal environment, driven by the concurrent enrichment of succinate-generating taxa and depletion of cross-feeding consumers that would otherwise convert it to protective SCFAs. The present multi-layer analysis provides a comprehensive mechanistic characterization of this process, situating succinate accumulation within a coordinated ecosystem reorganization whose carcinogenic consequences are evidenced simultaneously at the phenotypic, microbiome, metabolomic, and network levels.

These findings imply that the prevailing prebiotic framing of inulin may need reconsideration with direct relevance to contemporary dietary patterns. Inulin is among the most widely used functional food additives and supplement ingredients in the modern food supply, added to processed foods as a prebiotic fiber and fat replacer, and increasingly consumed in concentrated supplement form, precisely the purified, rapidly fermentable delivery format our findings identify as most consequential. Its growing presence in Western diets coincides with the sustained rise in early-onset colorectal cancer, a trend whose dietary determinants remain incompletely understood. While *Bifidobacterium* expansion is widely cited as a primary mechanism of inulin’s prebiotic benefit, the present findings demonstrate that this expansion occurs within a broader ecological context of depleted SCFA producers, disrupted cross-feeding networks, and accumulated fermentation intermediates, a context in which purported benefits are absent and carcinogenic vulnerability is increased. Inulin supplementation preconditions the gut microbiome toward a state of functional vulnerability, characterized by disrupted fermentation throughput and reprogrammed amino acid metabolism, that is amplified by carcinogen exposure into a tumor-promoting ecosystem. Future studies employing germ-free colonization models or targeted metabolite interventions will be important to establish the causal contributions of individual microbial taxa and metabolites identified here, and human cohort studies examining associations between inulin consumption, gut microbiome composition, and colorectal cancer risk will be essential to assess the translational relevance of these findings. Together, these results reframe inulin not as a universally beneficial prebiotic but as a dietary component whose consequences for gut ecosystem function and carcinogenic susceptibility depend critically on the biological context in which it is consumed.

## 5. MATERIALS AND METHODS

### Animal model and diets

Inbred wild-type male C57BL/6J mice were housed under temperature- and humidity-controlled conditions with *ad libitum* access to food and water. Mice were co-housed at 3–5 animals per cage. At 4 weeks of age, mice were randomly assigned to one of several compositionally defined diets containing 2.5% (w/w) cellulose and supplemented with an additional 7.5% (w/w) of a test fiber: cellulose (Cell), pectin, agar, or inulin. The control group received a diet containing only 2.5% (w/w) cellulose (Research Diets, Inc., New Brunswick, NJ). Detailed diet compositions are provided in **Table-S4**. Mice inclusion, randomization and effect sizes were determined as per prior 2defined criteria (46).

After a 4-week dietary adaptation period, mice within each dietary group were further subdivided into carcinogen-treated and baseline groups and maintained on their assigned diets for an additional 24 weeks. Mice in the carcinogen-treated groups received intraperitoneal injections of azoxymethane (AOM) at 7.5 mg/kg body weight, whereas baseline groups remained on their assigned diets without AOM administration.

The injection regiment was established based on pilot studies. Nine weekly AOM injections produced a high incidence of colorectal cancer irrespective of dietary treatment, whereas six injections failed to induce tumor formation. Therefore, an eight-injection protocol was selected for the present study. For the main experiment, mice assigned to the AOM groups received six weekly injections, followed by a 1-week interval before the seventh injection. The eighth injection was administered after an additional 2-week interval. Body weight was monitored weekly beginning with the first injection to assess overall health status throughout the experiment. Fecal samples were collected at defined timepoints from the beginning of the first AOM injection in carcinogen-treated groups. All animal procedures were approved by the Institutional Animal Care and Use Committee of Pennsylvania State University, under the protocol number PROTO202102016.

### Euthanasia and sample collection

Fecal samples were collected at specified intervals beginning with the first AOM injection in carcinogen-treated groups. At the study endpoint, mice were euthanized by carbon dioxide (CO₂) asphyxiation. CO₂ was supplied from a compressed gas cylinder fitted with a pressure-reducing regulator and an in-line flow meter and delivered through tubing to a dedicated induction chamber. Mice were euthanized individually in a euthanasia chamber that was not precharged with CO₂; gas flow was then initiated at a specified rate corresponding to a defined percentage of the chamber volume displaced per minute. Animals remained undisturbed in the chamber, with flow maintained for at least 1 min after respiratory arrest. Death was confirmed in every animal by absence of respiration and heartbeat together with loss of the pedal withdrawal reflex. These procedures were performed in accordance with the American Veterinary Medical Association *Guidelines for the Euthanasia of Animals*, under a protocol approved by the Institutional Animal Care and Use Committee of The Pennsylvania State University (PROTO202102016). Portal vein blood was collected into serum separator tubes (Becton Dickinson) and centrifuged to isolate hemolysis-free serum. Colonic tissues for protein extraction were flushed with ice-cold PBS and snap-frozen on dry ice or in liquid nitrogen. Tissues for RNA extraction were stored in RNAlater (Sigma-Aldrich) at −20 °C. Proximal-to-distal colonic Swiss rolls were fixed in neutral-buffered formalin for 24 h, transferred to 70% ethanol, and submitted to the Penn State Animal Diagnostic Laboratory for paraffin embedding, sectioning, and H&E staining. All other biological samples were stored at −80 °C until further analysis.

### Histological analysis

Colons were collected at euthanasia and fixed in 10% neutral buffered formalin for 24–48 hours. Tissues were then transferred to 70% ethanol and stored until paraffin embedding. Paraffin blocks were prepared by the Animal Diagnostic Laboratory at Pennsylvania State University. Sections of 5 μm thickness were cut from paraffin-embedded tissues and stored until staining. Before staining, sections were deparaffinized using an automated stainer (Leica). Slides were sequentially immersed three times in xylene, followed by 100%, 95%, 70%, and 50% ethanol, and finally tris-buffered saline, for 3 min at each step.

#### Hematoxylin and eosin (H&E) staining

Deparaffinized sections were stained with H&E by the Animal Diagnostic Laboratory at Pennsylvania State University.

#### Alcian blue staining

Alcian blue staining was performed on deparaffinized sections using a commercial kit (Vector Laboratories) as described in our earlier work. Briefly, sections were incubated in acetic acid solution for 3 min, followed by staining with Alcian blue solution for 30 min at room temperature. Excess stain was removed with acetic acid solution, and slides were rinsed under running water for 4 min. Nuclei were then counterstained with Nuclear Fast Red for 5 min and washed under running water for an additional 4 min. Finally, sections were dehydrated through a graded ethanol series (50%, 70%, 90%, and 100%) for 3 min at each concentration and mounted with DPX (Sigma).

### Image acquisition and analysis

The stained sections were imaged by DMi8 Leica microscope. The images were analyzed by ImageJ with Fiji extension.

### Amplicon sequencing

Approximately 50 mg of fecal material was used for microbial genomic DNA extraction using the ZymoBIOMICS 96 MagBead DNA kit (Zymo Research, Irvine, CA - D4308). Samples were randomly loaded into lysing tubes containing mixed-size zirconia beads (0.01–0.1 mm) and 750 μL ZymoBIOMICS lysis solution. The mixtures were homogenized using a FastPrep-96 instrument for 5 minutes and centrifuged at 10,000 × g for 5 min. DNA extraction was subsequently performed according to the manufacturer’s instructions.

Amplicon sequencing targeting the V4 region of the 16S rRNA gene was performed as previously described by Gohl et al. (47). In brief, the V4 region was amplified using primers 515F (GTGYCAGCMGCCGCGGTAA) and 806R (GGACTACNVGGGTWTCTAAT) primers), which included partial overhang sequences for i7 and i5 adapter addition. PCR products were diluted based on preliminary SYBR Green-monitored serial dilution trials (10× dilution). Diluted amplicons were indexed using dual 12-nt barcodes and quantified using PicoGreen (Life Technologies). Samples were then pooled at equimolar concentrations, gel-purified, and sequenced on an Illumina MiSeq using V3 600 cycle reagents run as 270x12x12x270.

### Microbiome data processing and analysis

Raw 16S rRNA gene sequencing data were generated across 4 independent sequencing runs and processed through QIIME2, generating amplicon sequence variant (ASV) tables, taxonomy assignments (GTDB-based), and representative sequences independently for each of four sequencing batches. Batch effects were assessed using Zymo mock community positive controls sequenced across all runs; ordination analyses (PCoA and NMDS) and PERMANOVA confirmed no significant technical batch effect (p = 0.364), and batch was therefore excluded as a covariate from downstream analyses. ASVs were collapsed to genus level using full GTDB lineage labels to avoid naming collisions, and count matrices were merged across batches. Following Callahan et al. (2016), genera present in fewer than 1% of samples and with mean relative abundance below 0.001% were removed, and sequences assigned to mitochondria, chloroplasts, or unclassified lineages were excluded (48). The final filtered dataset was split into AOM-treated and carcinogen-naive BASAL phyloseq objects for independent downstream analysis (49). Community-level composition was assessed by Bray-Curtis dissimilarity-based NMDS ordination and PERMANOVA (adonis2, 999 permutations, by = "margin") testing marginal contributions of diet, timepoint, and interaction, with homogeneity of dispersion evaluated by betadisper. For taxon-level analysis, a pseudocount of 1 was added to all counts prior to centered log-ratio (CLR) transformation, as count-ratio-based zero imputation resulted in removal of biologically relevant genera. CLR-transformed abundances were modeled per genus using a linear mixed model with a natural cubic spline for time:

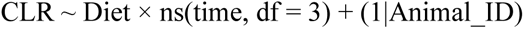

where time was treated as a continuous numeric variable (weeks 0, 6, 12, 14, 18, 19, 24), Diet as a categorical fixed effect with Control as the reference, and Animal_ID as a random intercept to account for repeated measures. Models were fitted by maximum likelihood and tested using ANOVA F-tests with Satterthwaite degrees of freedom approximation (lmerTest) (50). Genera with significant Diet × time interactions (p-value < 0.05) were retained for downstream interpretation. ICD-specific signatures were identified as taxa showing significant beta-coefficient divergence from the combined low-incidence fiber group (AgD, CCD, PCD) at four or more of seven sampling timepoints, with ICD exhibiting a uniquely extreme predicted abundance relative to all fiber diets at one or more timepoints. Model-predicted CLR trajectories and ICD-versus-fiber-mean beta-coefficient contrasts with 95% confidence intervals were generated for all significant taxa using the model’s own spline knot positions to ensure prediction consistency.

### 1H NMR spectroscopy

Hydrophilic metabolomic profiles of cecal contents were analyzed using proton nuclear magnetic resonance (¹H NMR) spectroscopy. Approximately 50 mg of cecal content was transferred into 2-mL screw-cap tubes. Ten to twenty silicate beads (0.1 mm) and 1 mL phosphate-buffered saline (PBS) prepared in 50% D2O containing 0.29 mM TMSP as an internal standard were added. Samples were homogenized at 4,260 rpm for two 30-s cycles with a 5-s pause between cycles using a Benchmark D2400 BeadBlaster Homogenizer. The homogenized slurry was then subjected to two freeze-thaw cycles using liquid nitrogen, followed by centrifugation at 17,000 × g for 15 min at 4 °C. Subsequently, 550 μL of supernatant was transferred into 5-mm NMR tubes for analysis using Bruker Avance NEO 600 MHz spectrometer equipped with a SampleJet sample changer (Bruker Biospin, Rheinstetten, Germany).

### Cecal metabolomics data processing and analysis

Cecal NMR spectra were processed and metabolite concentrations quantified as absolute values (µmol/g). For correlation-based network analyses, concentrations were subjected to CLR transformation followed by Pareto scaling (51). For differential abundance testing, raw absolute concentrations were used directly to preserve biological interpretability. Overall metabolome composition was assessed by PCoA on Bray-Curtis dissimilarity matrices with PERMANOVA (Diet × Treatment, 999 permutations). Between-group differences in individual metabolite concentrations were assessed using Welch’s two-sample t-test applied uniformly across all comparisons. Five comparisons were performed: ICD versus Control and ICD versus combined low-incidence fibers (AgD/CCD/PCD) within AOM mice; ICD versus Control and ICD versus combined low-incidence fibers within BASAL mice; and AOM versus BASAL within ICD mice. Effect sizes were computed as Cohen’s d. Metabolites significant at p-value < 0.05 in at least one comparison were considered differentially abundant.

### Multi-omics network integration

Spearman rank correlations were computed between CLR-transformed genus-level microbiome abundances and CLR + Pareto-scaled cecal NMR metabolite concentrations at week 24, independently for AOM-treated (n = 39) and BASAL (n = 22) cohorts. Features with zero variance were excluded prior to correlation testing, and p-value correction was applied across all pairwise tests within each cohort using the Benjamini-Hochberg method. Co-occurrence networks were constructed retaining edges with |ρ| > 0.449 and p-value < 0.1 for AOM, and |ρ| > 0.611 and p-value < 0.1 for BASAL, with thresholds selected empirically to maximize cluster significance while maintaining interpretable network density. Edge weights were defined as the Spearman correlation coefficient, and network files were exported for visualization and community detection in Cytoscape (52). Community detection was performed using ClusterOne (minimum cluster size = 5, minimum internal density = 0.3, edge weight = absolute correlation coefficient), with cluster significance assessed by permutation test, and cluster robustness was evaluated by Jaccard similarity-based bootstrap stability analysis comparing primary clusters against networks constructed at three progressively relaxed correlation thresholds (AOM: |ρ| > 0.40, 0.35, 0.30 without p-value correction; BASAL: |ρ| > 0.55, 0.50, 0.45 without p-value correction) (25). Group-specific bootstrap thresholds were applied to account for differences in network size and density between cohorts. Clusters were classified as Strong (Jaccard ≥ 0.3 at all three tiers), Moderate (Jaccard ≥ 0.3 at two tiers), Weak (one tier), or Unstable (none). Detailed breakdown of the cluster classification is provided elsewhere (**Table-S3**).

### Statistical analysis

All phenotype data are presented as mean ± SEM. Given the small sample sizes, data were analyzed using non-parametric statistical methods to minimize reliance on distributional assumptions, as done before (53). For comparisons involving more than two groups, Kruskal–Wallis tests followed by Dunn’s post hoc tests with Benjamini–Hochberg (BH) correction were used; for body-weight measurements, normality was additionally assessed using the Shapiro–Wilk test. For comparisons between two groups, the Wilcoxon rank-sum test was used. Statistical significance was defined as p-value < 0.05. For omics data, multiple comparisons were adjusted using BH false discovery rate. All analyses were performed in R (54) and Cytoscpae (52).

## Data and code availability

All the scripts used for the analyses are accessible via https://github.com/sumuduplus/AiW_project. Microbiome data have been deposited in the BioProject database under accession PRJNA1531042. Metabolomics data could be accessed via https://doi.org/10.5281/zenodo.22712212.

## Contributors

VS conceived and supervised the project, designed the study and led the conceptualization of the work. ST conducted the mouse experiments, supported by GY and UKG. ST, GY and UKG acquired the mouse experimental data. SR and ST performed the preliminary analysis, supported by TC and WG. SR performed data curation, formal analysis, validation and visualization, supported by JEB, JS and VS. FH, JEB and ADP additionally supported curation of the microbiome–metabolome data. JEB, JS, SRB, AKT and ADP contributed to the interpretation of the data. SR wrote the original draft of the manuscript, supported by ST and VS. VS was responsible for project administration, resources, supervision and funding acquisition, supported by JEB, JS, SRB, AKT and ADP. All authors critically reviewed the manuscript and approved the final version for publication.

*^#^ST and SR contributed equally*.

## Acknowledgement

The co-authors would like to acknowledge the Huck Institutes’ Genomics Core Facility (RRID:SCR_023645 and RRID: SCR_024530).

