## Supplementary material for "Dietary fiber-type-specific gut microbiome preconditioning drives differential susceptibility to sporadic colon cancer development": Figure S1-7 (from another version of this preprint: version 1, 2026-09-22)

Supplementary figure 1: Representative gross photographs of colons of the AOM cohort.

Supplementary figure 2: Representative gross photographs of colons of the basal cohort.

Supplementary figure 3: NMDS ordinations of AOM microbiome samples.

Supplementary figure 4-6: Predicted temporal abundance and differential beta-coefficient fluctuation overtime, for ICD significant genera identified (ordered by FDR adjusted p-value).

Supplementary figure 7: PCoA ordination of cecal metabolites across both cohorts.

Supplementary table 1: 72 significant taxa identified by the linear mixed model analysis.

Supplementary table 2: Differential metabolomics results.

Supplementary table 3: Stability specifications for microbiome-metabolome hubs identified.

Supplementary table 4: Breakdown of the diet compositions used.

**Supplementary Figures**

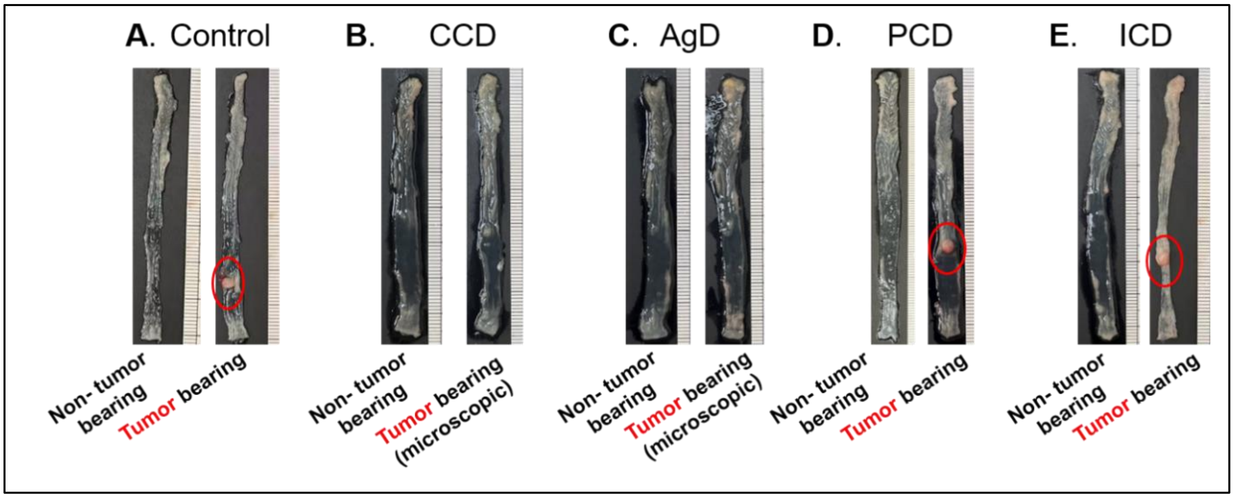


**Figure S1:** Representative gross photographs of colons (opened longitudinally) from control and DF intervention groups received AOM interjections. **A**. Control. **B**. Cellulose diet (CCD). **C**. Agar diet (AgD). **D**. Pectin diet (PCD). **E**. Inulin diet (ICD).

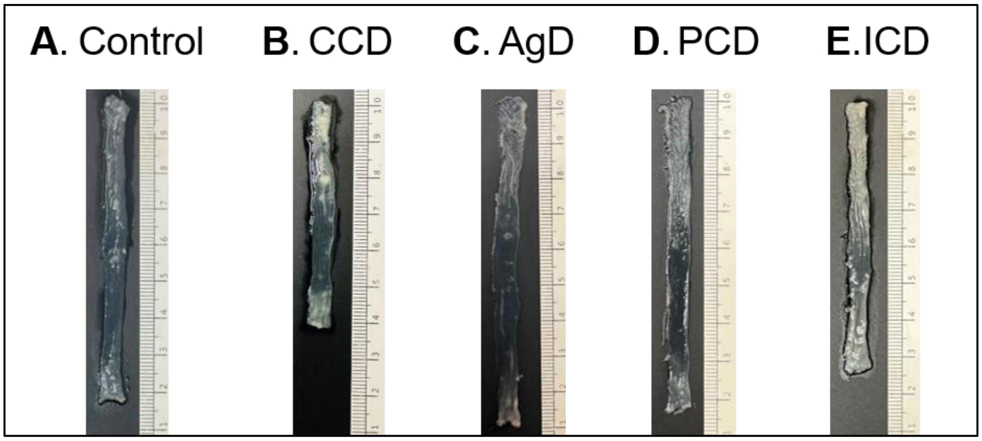


**Figure S2:** Representative pictures of colons (opened longitudinally) from DF intervention only groups (no AOM injection). A. Control. B. Cellulose diet (CCD). C. Agar diet (AgD). D. Pectin diet (PCD). E. Inulin diet (ICD).

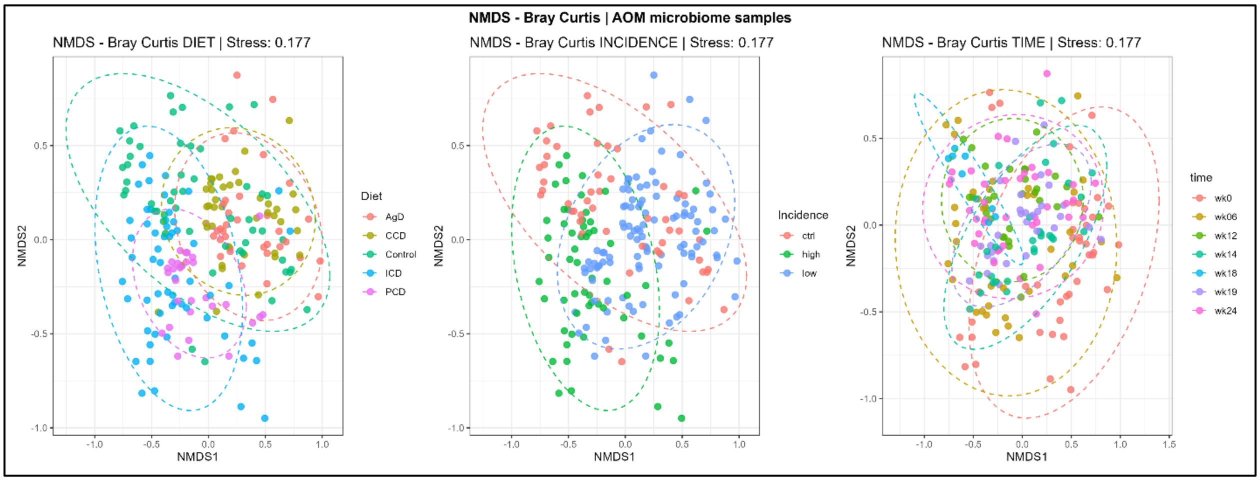


**Figure S3**: NMDS ordinations of AOM microbiome samples. Diets corresponding to Cellulose (CCD), Agar (AgD), Pectin (PCD), Inulin (ICD), and Control (ctrl). Time is denoted in weeks.

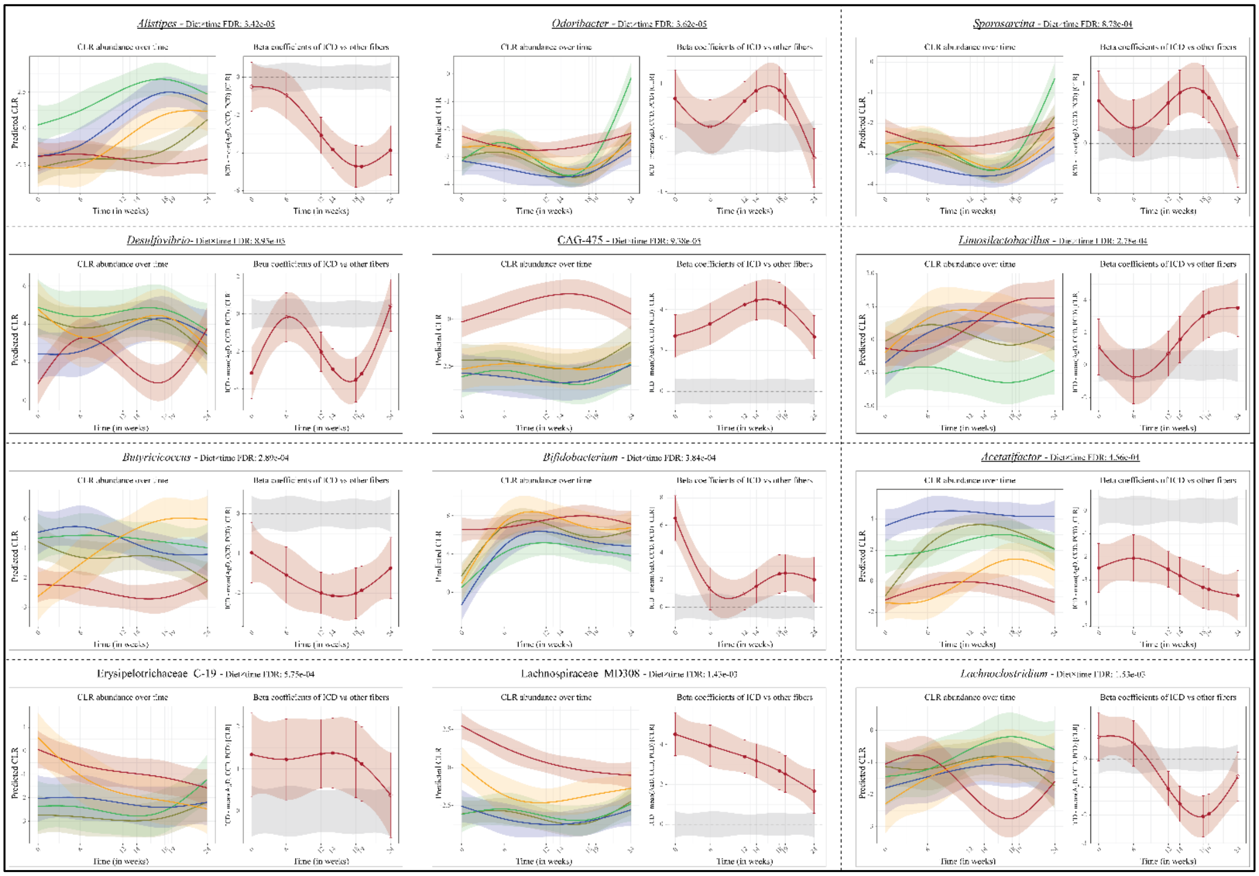


**Figure S4**: Predicted temporal abundance and differential beta-coefficient fluctuation overtime, for ICD significant genera identified: taxa 1- 12 (ordered by p-value). For each taxon, predicted CLR temporal abundance is plotted on the left and beta-coefficient shift of ICD, relative to collective fiber-groups, is plotted on the right. Significant deviations of beta-coefficients are denoted by the timepoint being filled. Standard deviation for plotted data is designated by the shaded range. Colors are as: Control (olive), Cellulose (blue), Pectin (yellow), Agar (green), Inulin (red), and collective fibers (grey).

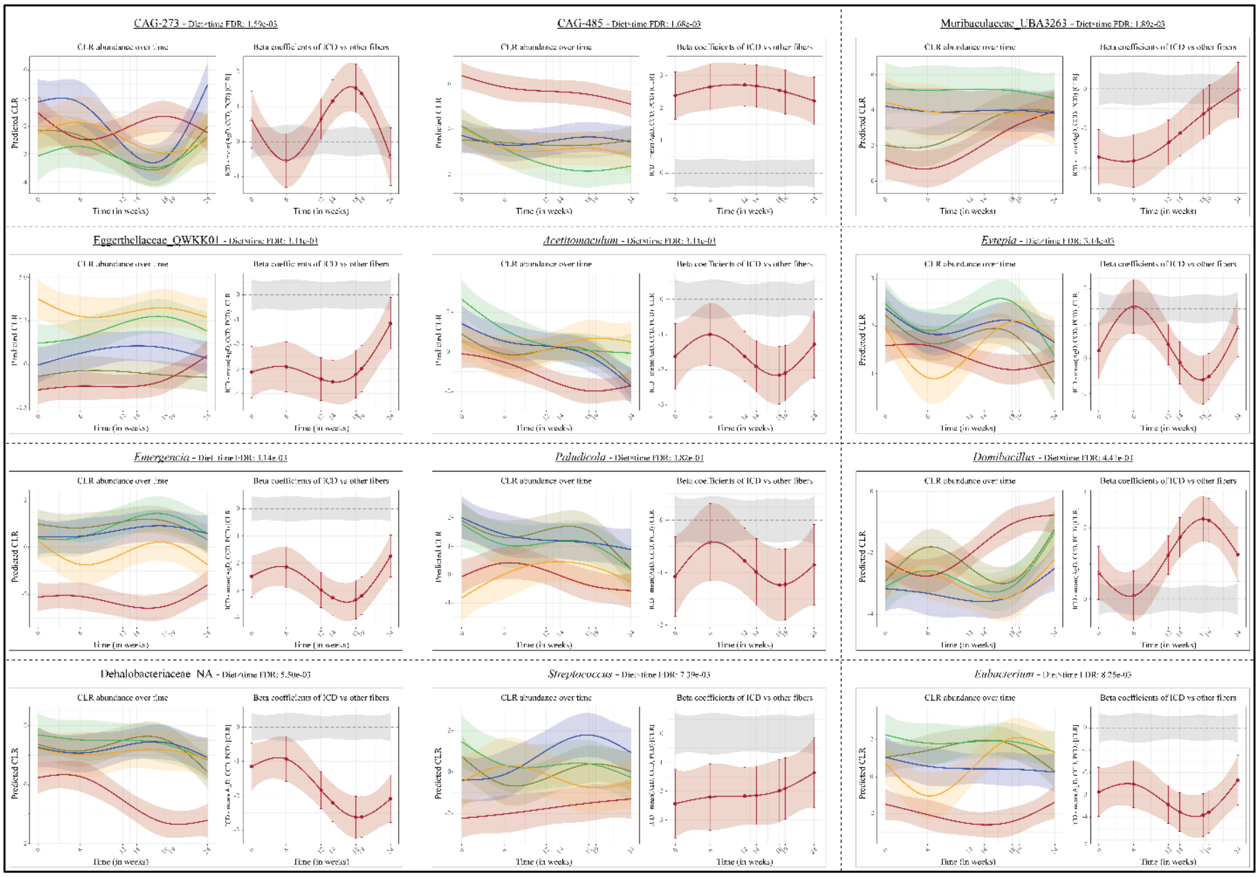


**Figure S5**: Predicted temporal abundance and differential beta-coefficient fluctuation overtime, for ICD significant genera identified: taxa 13- 24 (ordered by p-value). For each taxon, predicted CLR temporal abundance is plotted on the left and beta-coefficient shift of ICD, relative to collective fiber-groups, is plotted on the right. Significant deviations of beta-coefficients are denoted by the timepoint being filled. Standard deviation for plotted data is designated by the shaded range. Colors are as: Control (olive), Cellulose (blue), Pectin (yellow), Agar (green), Inulin (red), and collective fibers (grey).

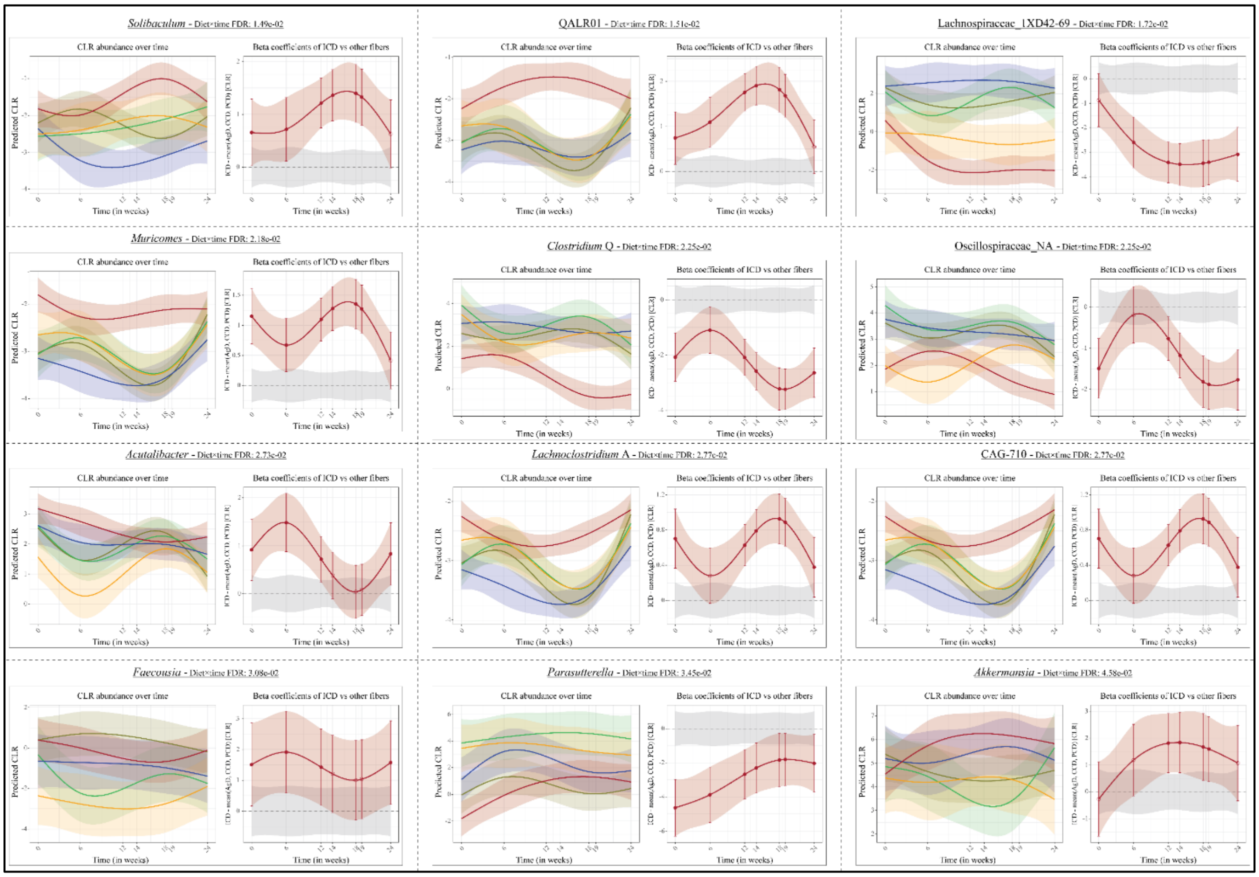


**Figure S6**: Predicted temporal abundance and differential beta-coefficient fluctuation overtime, for ICD significant genera identified: taxa 25- 36 (ordered by p-value). For each taxon, predicted CLR temporal abundance is plotted on the left and beta-coefficient shift of ICD, relative to collective fiber-groups, is plotted on the right. Significant deviations of beta-coefficients are denoted by the timepoint being filled. Standard deviation for plotted data is designated by the shaded range. Colors are as: Control (olive), Cellulose (blue), Pectin (yellow), Agar (green), Inulin (red), and collective fibers (grey).

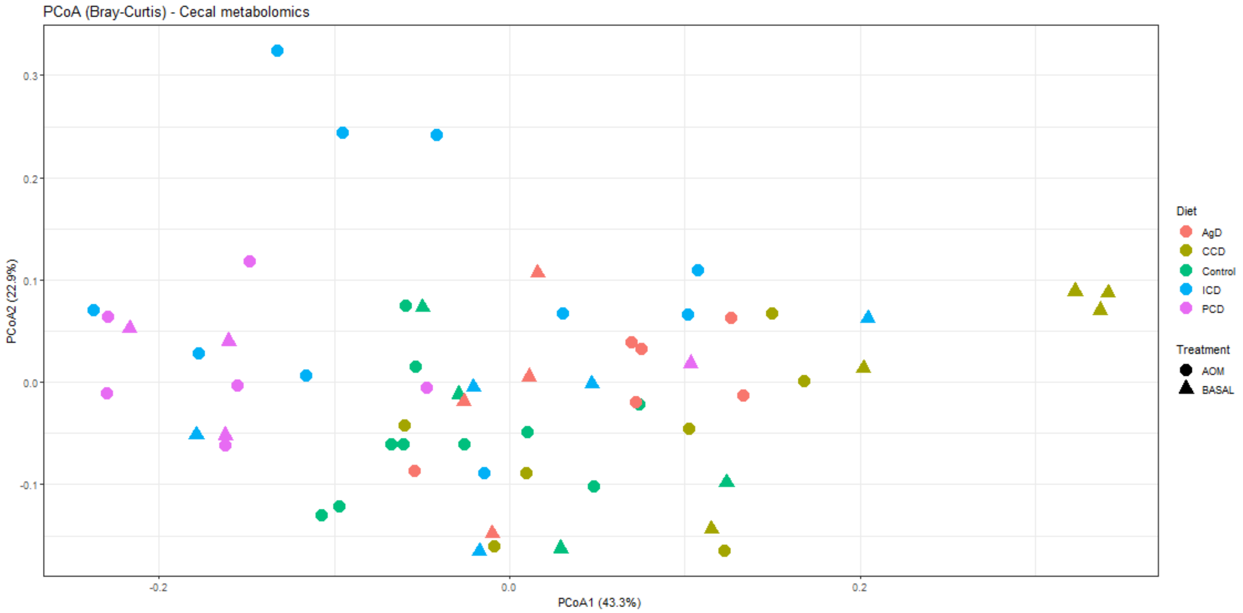


**Figure S7**: <u>PCoA ordination of cecal metabolites across all samples.</u> Bray-Curtis distance based Principal Coordinate Analayais of NMR cecal metabolomics data. Two cohorts used in the study are denoted by shape and diets are denoted by colors, as per the legend. Cellulose (CCD), Agar (AgD), Pectin (PCD), Inulin (ICD), and Control.

### Supplementary Tables

**Table S1**: <u>Significant taxa identified by the linear mixed model analysis.</u> Provided as "*Supplementary table 1 - MM_sig_diettime.csv*"

**Table S2:** <u>Differential metabolomics.</u> Provided as "*Supplementary table 2 -Differential metabolomics.csv*"

**Table S3:** <u>Stability specifications for microbiome-metabolome hubs identified.</u> Provided as *"Supplementary table 3 - Jaccard_stability_results.csv"*

**Table S4:** <u>Breakdown of the diet compositions used.</u> Provided as *"Supplementary table 4 - Diet formulations.xlsx*"
